# Evolution of transmissible spongiform encephalopathy and the prion protein gene (*PRNP*) in mammals

**DOI:** 10.1101/2020.01.27.920942

**Authors:** Brittaney L. Buchanan, Robert M. Zink

## Abstract

Wildlife managers are concerned with transmissible spongiform encephalopathies (TSEs) as they are currently incurable, always fatal, and have the potential to cross species boundaries. Although a wide range of mammals exhibit TSEs, it is currently unclear whether they are evolutionarily clustered or if TSE+ species are randomly distributed phylogenetically. We tested whether mammalian species with TSEs are phylogenetically underdispersed on a tree derived from 102 PRNP sequences obtained from the Orthologous Mammalian Markers database. We determined that the PRNP tree was topologically congruent with a species tree for these same 102 taxa constructed from 20 aligned gene sequences, excluding the PRNP sequence. Searches in Google Scholar were done to determine whether a species is known to have expressed a TSE. TSEs were present in a variety of orders excluding Chiroptera, Eulipotyphyla, and Lagomorpha and no marine mammals (Artiodactyla) were recorded to have a TSE. We calculated the phylogenetic signal of binary traits (D-Value) to infer if the phylogenetic distribution of TSEs are conserved or dispersed. The occurrence of TSEs in both trees is non-random (Species tree D-value = 0.291; PRNP tree D-value = 0.273), and appears to have arisen independently in the recent history of different mammalian groups. Our findings suggest that the evolution of TSEs develops in groups of species irrespective of PRNP genotype. The evolution of TSEs merits continued exploration at a more in-depth phylogenetic level, as well as the search for genetic combinations that might underlie TSE diseases.

## Introduction

Wildlife managers are concerned with transmissible spongiform encephalopathies (TSEs) as they are currently incurable, always fatal, and have the potential to cross species boundaries. Known TSEs include chronic wasting disease (CWD) in cervids, scrapie in sheep, bovine spongiform encephalopathy (BSE, also known as mad cow disease), transmissible mink encephalopathy (TME), feline spongiform encephalopathy (FSE) and Creutzfeld-Jacob in humans [1–3]. In response to health concerns of livestock and humans, research has focused on learning how species contract TSEs, how they are spread, causes of immunity, and prevention or cures [4].

Most researchers accept the hypothesis that resistance to TSEs in mammals results from certain genotypes found at the highly conserved prion protein gene (PRNP)[5]. TSEs are thought to be caused by the misfolding of the host’s prion protein (*PrP*) whose primary physiological function is not entirely clear. When correctly folded the prion protein has been theorized to localize at synaptic membranes and be related to normal synaptic functioning, signal transduction, and copper binding [6–9]. When misfolded the protein induces other prion proteins to misfold as well, followed by ultimately fatal accumulation in the central nervous system within the host [1, 5, 10]. Misfolded prion proteins can be spontaneously generated [4, 11] or introduced to the host by inoculation from the environment or through direct contact with infected individuals [5, 12]. Differences in mammalian prion proteins might reduce transmissibility between species because they function as a species barrier [5, 13].

A wide range of mammalian species exhibit TSEs, and it is currently unclear whether they are evolutionarily clustered, or whether TSE+ species are randomly distributed phylogenetically. If a species barrier inhibits horizontal transfer of TSEs, one might predict that related species would exhibit greater susceptibility to TSE expression. The reasoning for this prediction is that phylogenetically more distant relatives would be less similar genetically and, therefore, less susceptible to horizontal (cross-species) transmission. A phylogenetic test involves constructing a tree from PRNP sequences and testing whether species with TSEs are phylogenetically underdispersed, or clumped within clades [14]. In addition, because the PRNP gene might be under strong selection, it is important to document that the PRNP tree was topologically congruent with one that was not constructed with PRNP data. If the topology of the two trees differ significantly, it would suggest that selection has constrained the evolution the PRNP gene. If the presence of TSEs is phylogenetically clustered, and the two trees are more similar than one would expect by chance, it can be inferred that some lineages are predisposed, perhaps by their genetics, to acquiring this class of diseases. If TSEs are phylogenetically dispersed, and the two trees are similar, it would suggest that factors other than shared history explain the distribution of TSEs in mammalian taxa. Therefore, we have two objectives: 1) Determine if a mammalian species tree and PRNP gene tree have similar topologies and, 2) Determine if the presence of TSEs are phylogenetically dispersed in a species tree.

## Materials and methods

We used 102 aligned mammal sequences (Table S1) obtained from the Orthologous Mammalian Markers database (OrthoMam)[15]. In all phylogenic analyses, the platypus (*Ornithorhynchus anatinus*) was used as the outgroup. To determine whether a species is known to have a TSE, searches in Google Scholar were done using the scientific and common name of species combined with “TSE”, “transmissible spongiform encephalopathy”, “prion disease”, “CWD”, “chronic wasting”, “BSE”, “bovine spongiform”, “FSE”, “feline spongiforme”, “MSE”, and “mink spongiform”. Specific prion diseases were included in our search to broaden our list of taxa (Table S1). Many species appeared to have ambiguous evidence for TSE presence (Table S1), and we conservatively scored them as absent. Some species known to express TSEs lacked gene sequences that would have permitted including them in the species tree (e.g., moose, *Alces alces*, caribou, *Rangifer tarandus*, elk, *Cervus canadensis*, mule deer *Odocoileus hemonius*).

In addition to analyzing the aligned sequences available on Orthamam, we computed alternative alignments on the nucleotide data. We aligned sequences three separate ways in MEGA X [16] the default MUSCLE settings, the default MUSCLE settings followed with the program Gblocks [17–18] under stringent conditions to eliminate poorly aligned positions, and running the available Orthomam alignments only through Gblocks. The results of performing these alignment methods are the same as using aligned Orthomam sequences as-is. In addition, we constructed a phylogenetic tree using sequences of amino acids to determine if particular protein structures were associated with TSE+ species.

### Phylogeny construction

To construct a species tree independent of the PRNP gene, we selected 20 aligned gene sequences of coding regions (Table S2) for 102 species of mammals spanning 20 orders, 58 families, and 85 genera, and for which evidence of TSE presence/absence was available (Table S1). Using the aligned nucleotide coding regions, partitioned (by gene) analyses were run using the Bayesian Evolutionary Analysis by Sampling Trees 2 (BEAST 2) package [19]. The sequences were analyzed using the best fit model (HKY + G) identified using MEGA X [16]. We ran the analyses for 75,000,000 generations while sampling every 5,000 chains under a strict clock model and Yule speciation model. The first 10% of sampled trees were discard as burn-in. Two independent runs were performed with these specifications, and log files were combined in LogCombiner [20] to address low Effective Sample Size (ESS) values of parameters. Resulting trees were re-rooted to the platypus and exported as Nexus and Newick files. The PRNP gene tree was constructed using the same procedures as the species tree, with the best fit model identified as TN93 + G + I. To construct the PRNP gene tree, analyses ran for 10,000,000 generations while sampling every 5,000 chains under a relaxed log normal clock model and Yule model, along with three independent runs that were combined in LogCombiner [20]. The phylogenetic analysis of amino acids residues, obtained from Orthomam, followed the same protocol as the two preceding analyses. The amino acid PRNP gene tree had the same specifications as the nucleotide species tree with the best fit model identified as JTT + G and had three independent runs that were combined.

We mapped the presence or absence of TSE on the two trees using stochastic character mapping [21], which samples character histories based on their posterior probability distribution; we reconstructed the ancestral states using the equal rates model (run in Program R; version 3.5.2, R Development Core Team, 2018, Code S4). All analyses were done using the R package *phytools* version 0.6-99 [22]. The function *cophylo* was used to compare the species and gene tree. We also calculated the phylogenetic signal of binary traits (D-value) to infer if the phylogenetic distribution of TSEs are conserved or dispersed [23]. D-values close to 0 are not randomly distributed and are conserved, where if the value is close to 1 then the presence of the state is considered randomly distributed on the tree.

## Results

### Basic genetic results

The number of aligned base pairs ranged from 324 (*Monodelphis domesticus*) to 783 (*Bos taurus*, *Bos mutus*, *Bison bison*), and the total alignment included 861 base pairs, of which 486 were variable. Of the 287 total amino acids (no stop codons were noted), 166 were variable. Nucleotide composition differed little between species with and without TSE (Table S3). No amino acid positions separated TSE+ from TSE-species.

### Species and PRNP trees

Most of the internal nodes in the species tree (Fig. 1) were well supported with posterior probabilities over 0.90, with a few exceptions close to the terminal tips (Fig 1). The topology is consistent with current taxonomy, at least to the extent that species from the same orders are supported as clades. In contrast, the PRNP gene tree has relatively few strongly supported nodes (Fig 1), although the topology is also consistent with current mammalian ordinal taxonomy. Both trees are topologically congruent, with most of the discrepancies occurring at poorly supported nodes deep in the trees (Fig 1). The tree constructed from amino acids (Fig 2) is congruent with both the species and PRNP trees.

**Fig 1.**
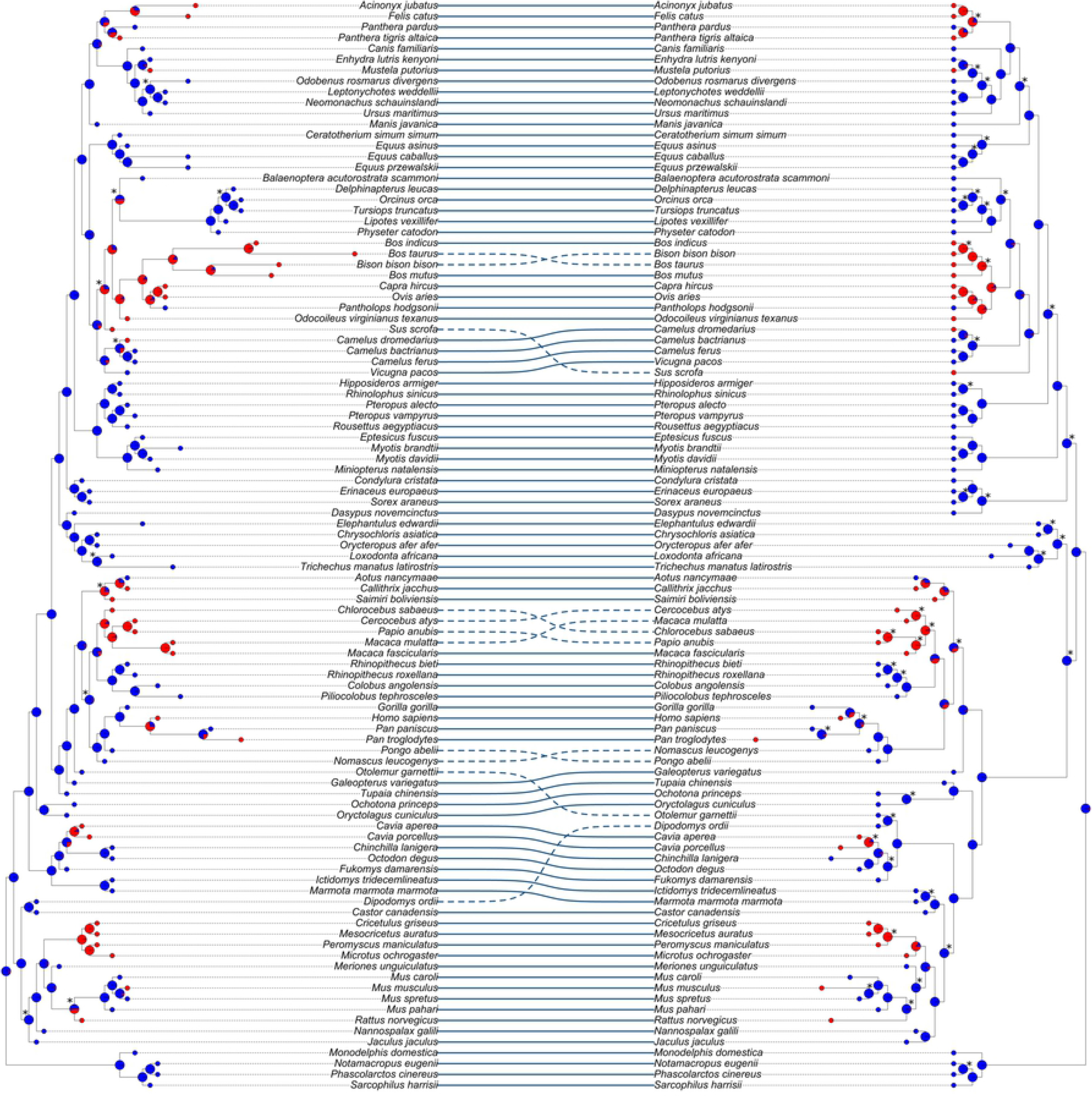
Species Tree (left) and PRNP Gene Tree (right) Comparison using Nucleotides. Compiled using 20 autosomal genes and rooted with platypus (excluded from figure). Positive TSE presence (red) and absence TSE (blue) shown at tips with orders that contain 2 or more species labeled. Posterior probabilities less than 0.90 and stochastic character mapping probabilities (as pies) displayed at nodes. Congruent tips are connected by solid lines, whereas topological differences between the trees are connected by dashed lines.

**Fig 2.**
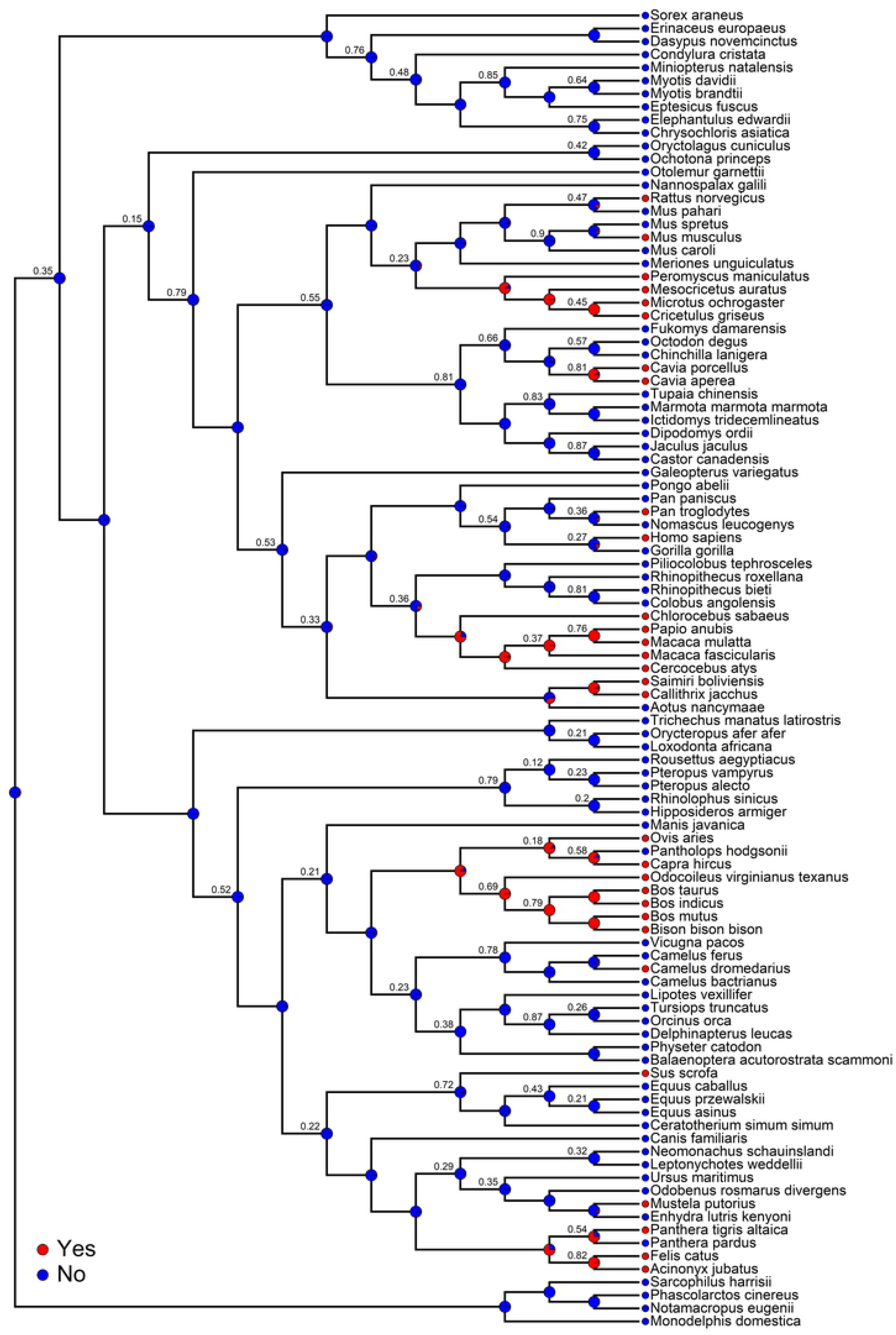
PRNP Gene Tree Created using Amino Acids. Compiled using the translated PRNP gene and rooted with platypus (excluded from figure). Positive TSE presence (red) and absence TSE (blue) shown at tips with orders that contain 2 or more species labeled. Posterior probabilities less than 0.90 and stochastic character mapping probabilities (as pies) displayed at nodes.

### Reconstruction of TSE evolution

Because of the congruence of the two trees, we focused on the results from the species tree. TSEs are present in a variety of orders excluding Chiroptera, Eulipotyphyla, and Lagomorpha. No marine mammals (Artiodactyla) have been recorded to have a TSE. According to the ancestral reconstruction, TSEs appear to have arisen relatively recently in TSE+ groups, with the basal condition being absence of TSEs. The reconstruction of TSE evolution is also notable in that there was only one hypothesized transition from TSE presence to absence (Tibetan antelope, *Pantholops hodgsonii*). The presence of TSEs is non-random (D-value = 0.291), suggesting that TSE presence is relatively conserved. Therefore, the distribution of TSEs is not randomly distributed across the phylogeny (Fig. 3). The results for the PRNP gene alone were identical to the results inferred from the species tree.

**Fig 3.**
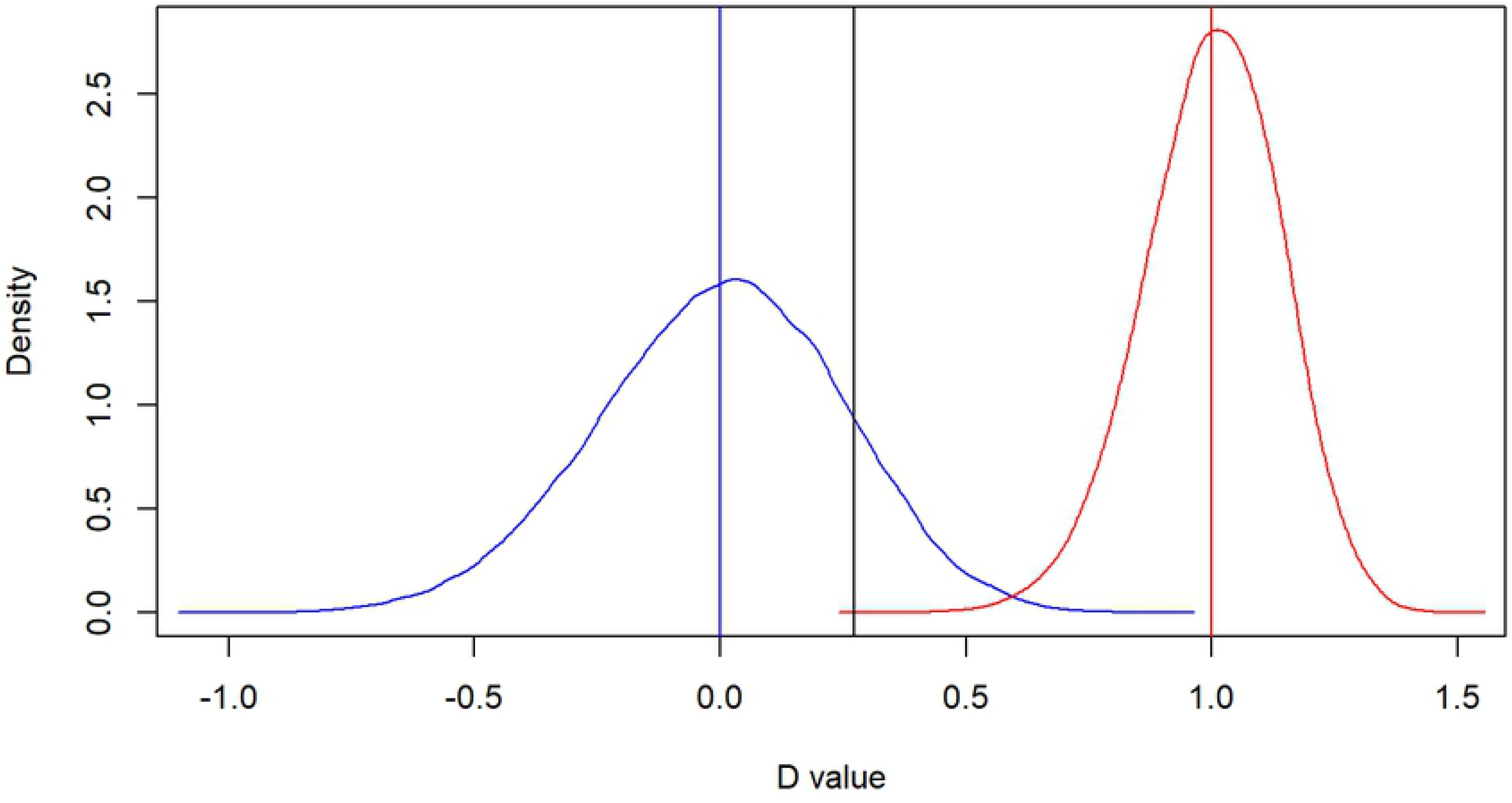
Density plot of scaled observed value of D for the species tree. The observed value of D for the species tree (D = 0.291) in black compared to simulated values of D = 0 (blue), representing the traits being phylogenetically conserved as expected under a Brownian threshold model (p = 0.115), and D = 1 (red) as the traits being phylogenetically random under a Brownian threshold (p = 0). PRNP tree has similar results (not shown) with observed value of D = 0.273. P = 0.135 for the simulated value of D = 0, and a p = 0 for the simulated value of D = 1.

## Discussion

Our species tree and the tree inferred solely from the PRNP gene (Fig. 1) closely match accepted mammalian phylogenetic trees [24–28]. Therefore, our species tree provides a glimpse into the evolution of TSEs. The occurrence of TSEs is not randomly distributed across the mammal phylogeny. TSEs appear to have arisen independently and recently in several major mammalian groups whereas they are absent in others; had information for the 20 genes been available, a group of cervid species (e.g., moose, caribou, elk), all which exhibit TSEs, would have been clustered with *Odocoileus virginianus*. The lack of amino acid substitutions unique to all TSE+ or TSE-species suggests there is not a particular set of amino acid substitutions that either provides resistance or susceptibility. In sheep amino acids positions relevant to scrapie resistance are 136 (A/V), 154 (R/H) and 171 (Q/R/H), with the 136A/154R/171R genotype conferring complete or nearly complete resistance [29]. We did not find this genotype in any other mammalian taxa. Our ancestral reconstructions included only one instance of a reversal from TSE presence to absence, suggesting that mutations conferring resistance are relatively rare. The nonrandom occurrence of TSEs in some mammalian orders (e.g., rodents, bovids, felines, cervids) suggest that TSEs are a recently evolved class of mammalian disease, which could explain why TSEs are nearly always fatal. Rongyan et al. (2008:650) suggested that “no dramatic sequence changes have occurred to avoid cross-species TSE infectivity.” Why TSEs are not more widespread across mammals is unclear at this time. It seems possible that the evolution of TSEs is independent of PRNP genotype.

Given the high fatality rates of TSEs, one might expect strong selection on the PRNP gene. Balancing selection in sheep [30] and strong purifying selection for PRNP CDS has been implicated in cattle [31]. In contrast, the congruence between trees reconstructed from nucleotides (Fig. 1) and amino acid residues (Fig. 2) suggests that selection has not yet played a major role in the evolution of the PRNP gene. That is, if there were a common PRNP genotype at the amino acid level that conferred resistance to TSEs, those species ought to have been grouped together on the amino acid tree in a way that conflicts with the species tree.

Reconstruction of TSE evolution suggests the ancestral state is the absence of TSEs (Fig. 1), and that certain orders of mammals are apparently at greater risk of developing or contracting these diseases. Alternatively, it is possible that our knowledge of the occurrence of TSEs is incomplete, and one interpretation of our analysis is that all mammalian orders are susceptible, which could be confirmed by more extensive testing. If TSEs are a relatively recent phenomenon in mammals, perhaps enough time has not passed for crossing of species-group barriers. Rongyan et al. (2008) noted that scrapie has been endemic in the United Kingdom for more than 200 years and yet has not crossed the species barrier into humans.

## Conclusions

Understanding how diseases have evolved plays a crucial part in determining which species are currently most at risk, and if the possibility for others to become at risk is an immediate concern. Our findings show that the evolution of TSE+ species is localized, non-random, and recently developed in groups of species irrespective of PRNP genotype. As the PRNP gene has been associated with varying susceptibility to TSE diseases in past studies [11, 32–34], future studies should focus on other genes. For example, in some cattle breeds and the gayal (*Bos frontalis*), a 23-bp deletion in the PRNP promoter region is associated with susceptibility to bovine spongiform encephalopathy (BSE) [35–36], although this was not found for white-tailed deer and mule deer (Zink et al. *in review*). Most research into TSEs has involved species such as cows, sheep, deer, rodents, and select primates, which could illuminate how TSEs could cross the species barrier into humans. However, as shown by our list of TSE+ species (Table S1) there are many species that are as yet unstudied. Therefore, our list of TSE+ species could be incomplete. The evolution of TSEs merits continued exploration at a more in-depth phylogenetic level, as well as the search for genetic combinations that might underlie TSE diseases.

## Acknowledgments

We thank Hernan Vazquez-Miranda for help in conceptualizing this project and advice on phylogenetic methods, along with Chris Chizinski and Jeffrey Lusk for their insight and helpful comments.

## Supporting information

**S1 Table. List of species with a record of contracting a TSE or not with related references**.

**S2 Table. List of genes used for BEAST analysis, all gene sequences obtained from Orthomam**.

**S3 Table: Nucleotide composition for the PRNP gene averaged by TSE+ and TSE-species**

**S4 Code: R Software (version 3.5.2) code used to visualize phylogenies and run analyses**

